# Developmental effects on pattern visual evoked potentials characterized by principal component analysis

**DOI:** 10.1101/2020.12.11.420158

**Authors:** Carlyn Patterson Gentile, Nabin R. Joshi, Kenneth J. Ciuffreda, Kristy B. Arbogast, Christina Master, Geoffrey K. Aguirre

## Abstract

**Purpose:** Peak amplitude and peak latency in the pattern reversal visual evoked potential (prVEP) vary with maturation. We considered that principal component analysis (PCA) may be used to describe age-related variation over the entire prVEP time course and provide a means of modeling and removing variation due to developmental age.

**Methods:** prVEP was recorded from 155 healthy subjects ages 11-19 years at two timepoints. We created a model of the prVEP by identifying principal components (PCs) that explained >95% of the variance in a “training” dataset of 40 subjects. We examined the ability of the PCs to explain variance in an age- and sex-matched “validation” dataset (n=40) and calculated the intra-subject reliability of the PC coefficients between the two timepoints. We explored the effect of subject age and sex upon the PC coefficients.

**Results:** Seven PCs accounted for 96.0% of the variability of the training dataset and 90.5% of the variability in the validation dataset with good within-subject reliability across timepoints (R>0.7 for all PCs). The PCA model revealed narrowing and amplitude reduction of the P100 peak with maturation, and a broader and smaller P100 peak in males compared to females.

**Conclusions:** PCA is a generalizable, reliable, and unbiased method of analyzing prVEP. The PCA model revealed changes across maturation and biological sex not fully described by standard peak analysis.

**Translational relevance:** We describe a novel application of PCA to characterize developmental changes of prVEP in youth that can be used to compare healthy and pathologic pediatric cohorts.

## INTRODUCTION

Visual evoked potentials (VEP) are a relatively inexpensive and easily implemented method of probing the cortical visual system. Pattern reversal (pr) VEPs, which are generated by viewing reversing checkerboard patterns, have the additional desirable property of relatively high intra-subject reliability in adults^1^. The prVEP can be helpful for clinical diagnosis and monitoring in demyelinating conditions such as multiple sclerosis^2^, identifying sources of visual acuity loss^3^, and shows differences in a broad range of neurologic conditions including concussion^4–7^ and migraine^8^.

Despite the high potential of VEP as a diagnostic tool, prVEP interpretation faces unique challenges in the pediatric population. Multiple studies have found the prVEP varies during maturation, effects of which can be seen into early adulthood^9–13^. Age variation may confound studies that compare healthy and pathologic groups, either because of unmatched age differences in the populations, or more subtly, from a non-specific effect of the disease upon maturation. This is especially true for longitudinal prVEP measurements. Few studies have examined the stability of VEP across time in the pediatric population. One study showed high intra-subject variability of flash VEP signals collected in children 10 months apart on average^14^. This variability was higher than adult studies^14^. The ability to define and account for developmental changes is critical for the interpretation of VEPs in the pediatric population.

Standard prVEP analysis relies on peak amplitude and peak latency measurements of predictable positive and negative peaks that occur in the first 150ms of the prVEP^15^. Prior studies of developmental effects upon the prVEP have found differences at multiple prVEP peaks as a function of age, although these results have been somewhat inconsistent. Some studies have found general decreases in peak latency^9,10^, while others have seen increases^13^, or no change^11^. Peak amplitude tends to decrease with increasing age^10–12^, however some studies found that this was not the case for all peaks studied^10^. Inconsistencies in these results may be due, in part, to the peaks analyzed, how peaks were defined, and differences in non-local temporal VEP waveform shape that are not fully captured by restricting analysis to pre-defined maxima and minima.

Principal component analysis (PCA) offers a potential solution to these challenges. PCA partitions the variability of a dataset into orthogonal, uncorrelated dimensions called principal components (PCs). As biologically relevant variation may be restricted to a subset of possible outcomes, PCA, like peak analysis, may be used to reduce the dimensionality of the VEP, but with the additional advantage of preserving informative variability of the global temporal signal^16^. This approach does not require specific reference to pre-defined, predictable VEP peaks. Further, once characterized by weights on a set of components, the effects of a continuous biological variable like developmental age can be modeled and accounted for in future analysis.

We used PCA to analyze prVEP timeseries data from a single active electrode collected from a large youth cohort.. Using separate training and validation datasets, we demonstrate that a particular set of PCA components provide for generalizable analysis of prVEP data. Using measurements from two separate sessions in the validation dataset, we further show that the coefficients measured for these components have good reproducibility for a given subject across a short time span (weeks to months). Finally, we use this method to characterize developmental differences in our pediatric cohort over a span of years.

## METHODS

### Subjects

Subjects between the ages of 11 and 19 years were recruited from a local Philadelphia-area school as part of a study on youth concussion conducted as part of the Children’s Hospital of Philadelphia Minds Matter Concussion Program. Specifically, subjects were recruited through the sports teams of their local school to collect pre-season VEP data on healthy athletes. Subjects with a past history of concussion were at least 30 days from their most recent concussion and had resolution of their concussion symptoms. Consent was obtained from subjects and guardians, and all studies were approved by the Children’s Hospital of Philadelphia Institutional Review Board and were in accordance with the Declaration of Helsinki. PrVEPs were collected on 155 subjects between February 2018 and February 2020. One-hundred and five subjects had at least two recorded sessions separated by 0.7 to 17 months. For 90 of the subjects the sessions were separated by <6 months. Subject demographics including age, sex, race, ethnicity, and past medical history (PMH) were recorded. Any medications subjects were using were recorded. As part of the broader study, all subjects filled out a post-concussion symptom inventory (PCSI). Subjects were prescreened to ensure normal or corrected normal visual acuity under binocular and monocular viewing conditions using the Snellen’s visual acuity chart at 10 feet (5 subjects wore contacts, and 9 subjects wore glasses). Those with corrective lenses wore them during testing.

### Study paradigm

#### Viewing conditions

Subjects were seated in a well-lit, quiet room approximately 1 meter distant from a visual stimulus monitor. The room was not darkened. The luminance of the white walls adjacent to the testing apparatus was 53.0 cd/m^2^ as measured using a spectrophotometer (PR670, Jadak Inc, North Syracuse, NY). Data were collected under binocular viewing conditions. Visual stimuli were presented on a monitor as part of a vision testing system (NOVA, Diopsys®, Pine Brook, NJ). The monitor was a 17” LCD screen with 1280 × 1024 resolution, and a 75 Hz refresh rate. The screen was factory calibrated with gamma correction. The measured luminance of the stimulus background (half-on primaries) was 103.0 cd/m^2^.

#### Visual stimulus and paradigm

Visual stimuli were ISCEV Standard large checks^15^ presented in a wide field 85% contrast checkerboard with a pattern reversal rate of 2 reversals per second. Checks were 0.97-degree visual angle. Subjects were instructed to fixate on a central red ‘x’ for the duration of the stimulus presentation. Stimuli were presented for a continuous 20 seconds for a total of 40 reversals in a single block. The block was repeated 5 times in a single session. There was approximately 30 to 60 seconds in between blocks.

#### VEP recordings

VEPs were recorded with a vision testing system (NOVA, Diopsys®, Pine Brook, NJ). Two reference electrodes were placed at Fz, and laterally to Fz at Fp1, with the active electrode placed at Oz following the 10-20 international EEG placement criteria^17^ based on Diopsys system recommendations. Data were recorded at a sampling rate of 1024 samples/second. Raw voltage-by-time data were exported for subsequent analysis. Visual stimulus presentation was time-locked to VEP recording.

### Data analysis

Preregistration for this study can be found at the following link: https://github.com/pattersongentilelab/preregistrations. Data analysis was performed using custom software written in a coding program (Matlab, MathWorks, Natick, MA).

#### VEP preprocessing

Notch filters were applied at 60 Hz and 120 Hz to remove powerline noise. Data were parsed into 500ms intervals corresponding to each pattern reversal. Pattern reversals were excluded if there was a voltage change of >1 mV indicating a large non-physiologic change in voltage (~2% of reversals were discarded by this criterion). Signal-to-noise ratios (SNR) were calculated for each block by taking the mean squared of each time point divided by the standard deviation squared, then taking the mean SNR across all time points. Blocks with an SNR of <0.03 (level determined by distribution of SNR values across all blocks) were excluded (3% of blocks). All VEP trials were adjusted to a baseline of 0 by subtracting the mean of the first 50ms.

#### VEPs

The mean prVEP for each subject was calculated across all reversals collected in a given session. Each session consisted of 5, 20-second blocks. Representations of prVEPs show the mean across-subject VEP with 95% confidence intervals obtained by bootstrap analysis across subjects with replacement.

#### PCA model

Twenty females and 20 males were randomly selected from a pool of 90 subjects who had at least two recorded sessions, less than 6 months apart (Figure S1). These subjects constituted the training dataset. Mean prVEP across all trials, and across the two sessions from the training dataset, were used for PCA calculations. PCA was calculated using singular value decomposition (SVD):

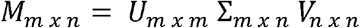

where Matrix M is a *m* × *n* matrix, Matrix U is a is a *m* × *m* orthogonal matrix, ∑ is a *m* × *n* rectangular diagonal matrix, and V is a *n* × *n* orthogonal matrix. For our dataset, matrix M consisted of m=40 subjects and n=512 voltage points spanning 500ms sampled at a rate of 1024 Hz. The PCA was decentered (i.e., the mean was not subtracted from the responses for the analysis). The initial PCs that described >95% of the total variability were retained for subsequent analysis. We refer below to this set of PCs as the “PCA model”.

#### PCA model validation for generalizability and retest reliability

Forty age- and sex-matched subjects were selected from the remaining pool. This validation dataset was projected onto the PCA model. The distribution of coefficients for each PC were compared between the training and validation datasets using a two-sample Kolmogorov-Smirnov (KS) test.

We examined the reproducibility of coefficients within subject. Each session from the validation dataset was projected on to the PCA model. We calculated the Pearson correlation coefficient between PC coefficients from session 1 and session 2 across subjects. We then calculated the distance between session 1 and session 2 within the 7 PC dimensions for all subjects to determine if a subject could be correctly matched between their first and second sessions. Rank order for the smallest Euclidean distance was determined for the subject’s own match between session 1 and session 2, compared to combinations with all other subjects.

#### Sex and age comparisons

Categorical variation across the subject group in sex, and continuous variation in age, were each modeled by an ANOVA for which the covariates were created from the 7 PC coefficients from each subject. We simulated prVEP signals for a 10-year old, 15-year old, and 20-year old subject (regressing out the effect of sex) by obtaining the PC coefficients of the 7 PCs for this age group and fitting to the PCA model.

#### Peak Analysis

The N75 and P100 peaks were defined by identifying the local minimum in the 60 – 90ms range, and the local maximum in the 90 – 130ms range, respectively. This was done from the mean VEP waveform for each session for the comparison across sessions, and from the mean VEP waveform across sessions for the age correlation. Pearson Correlation Coefficients between session 1 to session 2, and correlation with age were calculated for these three variables.

## RESULTS

### Seven components are sufficient to capture prVEP variability

The prVEP (averaged across sessions) from 40 randomly selected subjects (20 females, 20 males) was used to derive the PCA model (i.e., training dataset). Specifically, PCA was used to capture the variability in the VEP waveform across these 40 subjects (Figure 1a). The mean prVEP for the training dataset is shown in Figure 2a. Seven PCs (Figure 2b) accounted for 96.0% of the variability in this sample (Figure 2c). We refer to these 7 PCs generated by the training dataset as the PCA model. As the PCA was decentered, the first PC approximates the mean prVEP. PCs 2 though 7 do not clearly map to particular peaks or points of time within the prVEP signal. However, certain peaks are represented in particular ways within these PCs. For instance, PC 2 has the effect of broadening the P100 peak. PC 3 has the effect of shortening the peak latency of P100 and increasing the P100 amplitude. PC 4 primarily broadens the P100 peak. These complex changes in peak shape would not be fully captured by traditional peak analysis techniques.

**Figure 1:**
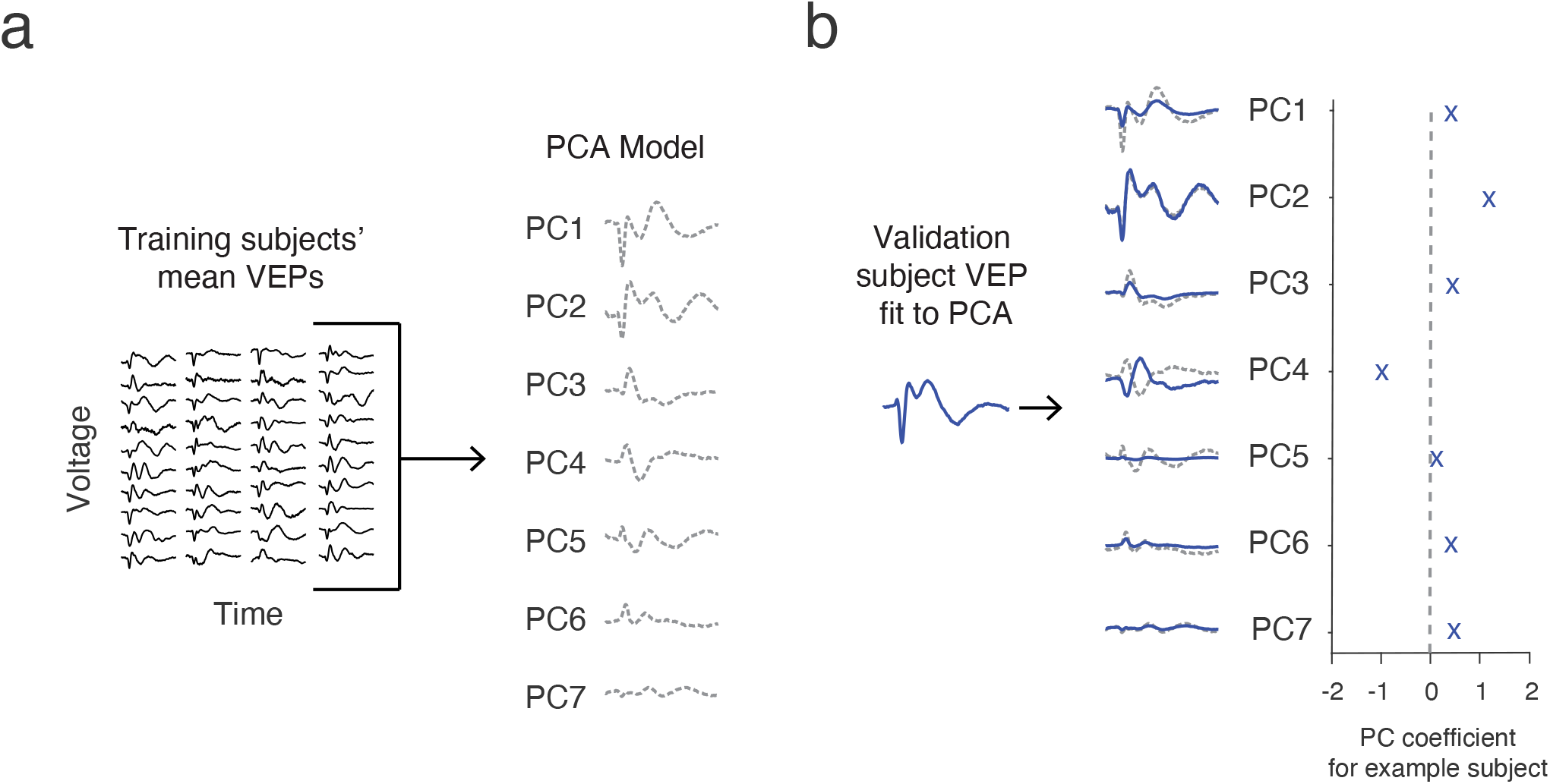
Schematic demonstrating generation and use of the PCA Model. (a) Forty randomly selected subjects were used to generate the PCA model. 7 PCs (gray dotted waveforms) capture 96.0% of the variability across these 40 subjects. (b) An example subject from the validation dataset is fit to the PCA model. This subject was not used to generate the original model. The example subject’s mean VEP waveform (blue) is fit to the PCA model (the PCs multiplied by the coefficients for the example subject are shown in blue lines overlapping the original PCs in dotted gray) and a PC coefficient is generated for this subject for each of the 7 PCs (blue “x”).

**Figure 2:**
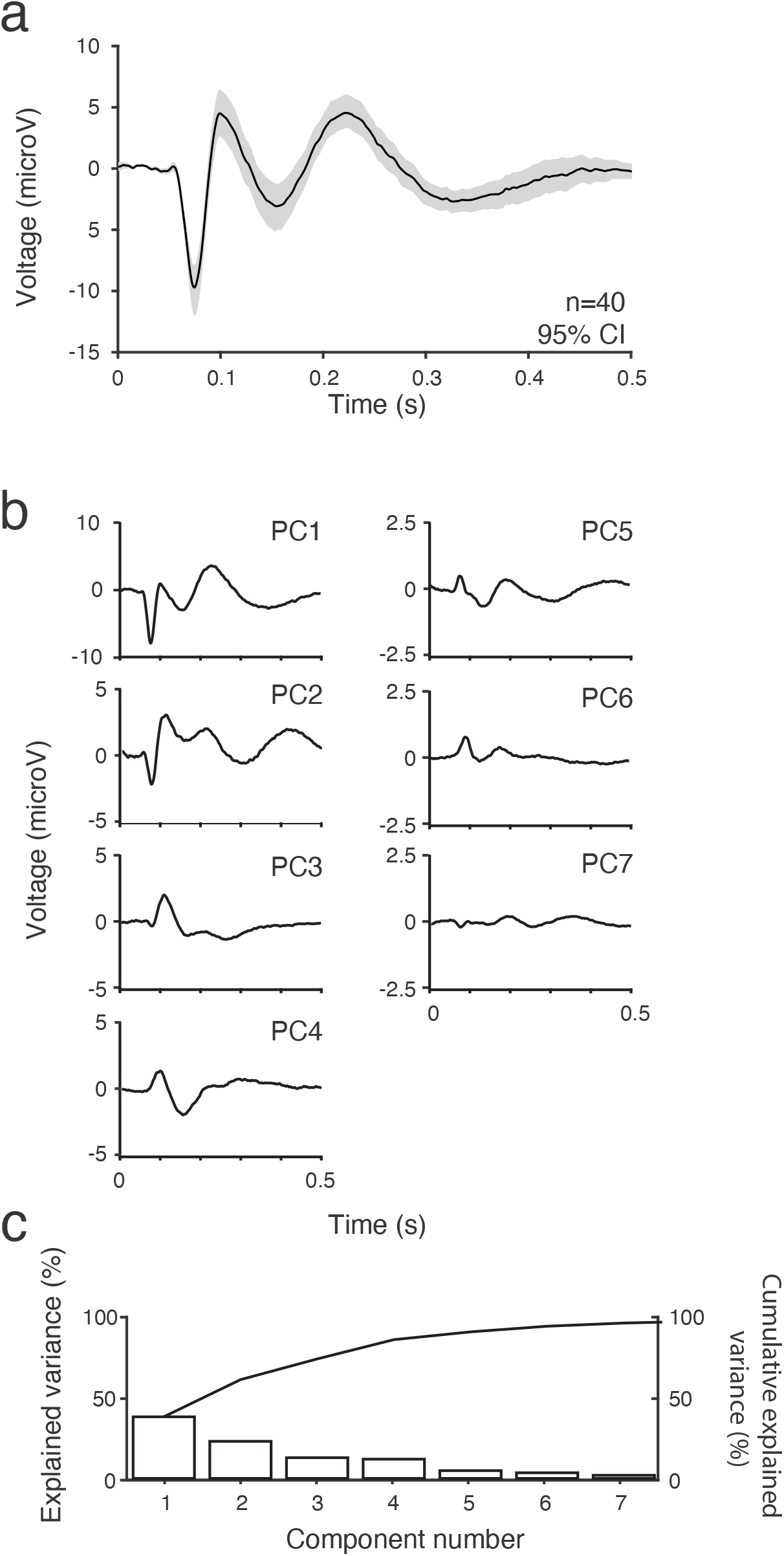
PCA Model. PCA was generated with 40 randomly selected subjects (20 males, 20 females). (a) mean VEP response across all 40 training subjects; gray represents 95% confidence interval (CI) by bootstrap analysis. (b) Principal components (PCs) 1 to 7, which account for 96.0% of the variability in the data. (c) Scree plot showing percent explained variance of PCs 1 to 7.

### The PCA model yields similar PC coefficients in an independent validation dataset

Forty age- and sex-matched subjects were selected to validation the generalizability of the PCA model. There were no significant differences in patient demographics including race/ethnicity and PMH between the training and validation subjects (Table 1). There was a significant difference of the number of subjects on stimulant medications for ADHD, with more subjects on these medications in the validation compared to the training group (Table 1).

**Table 1:**
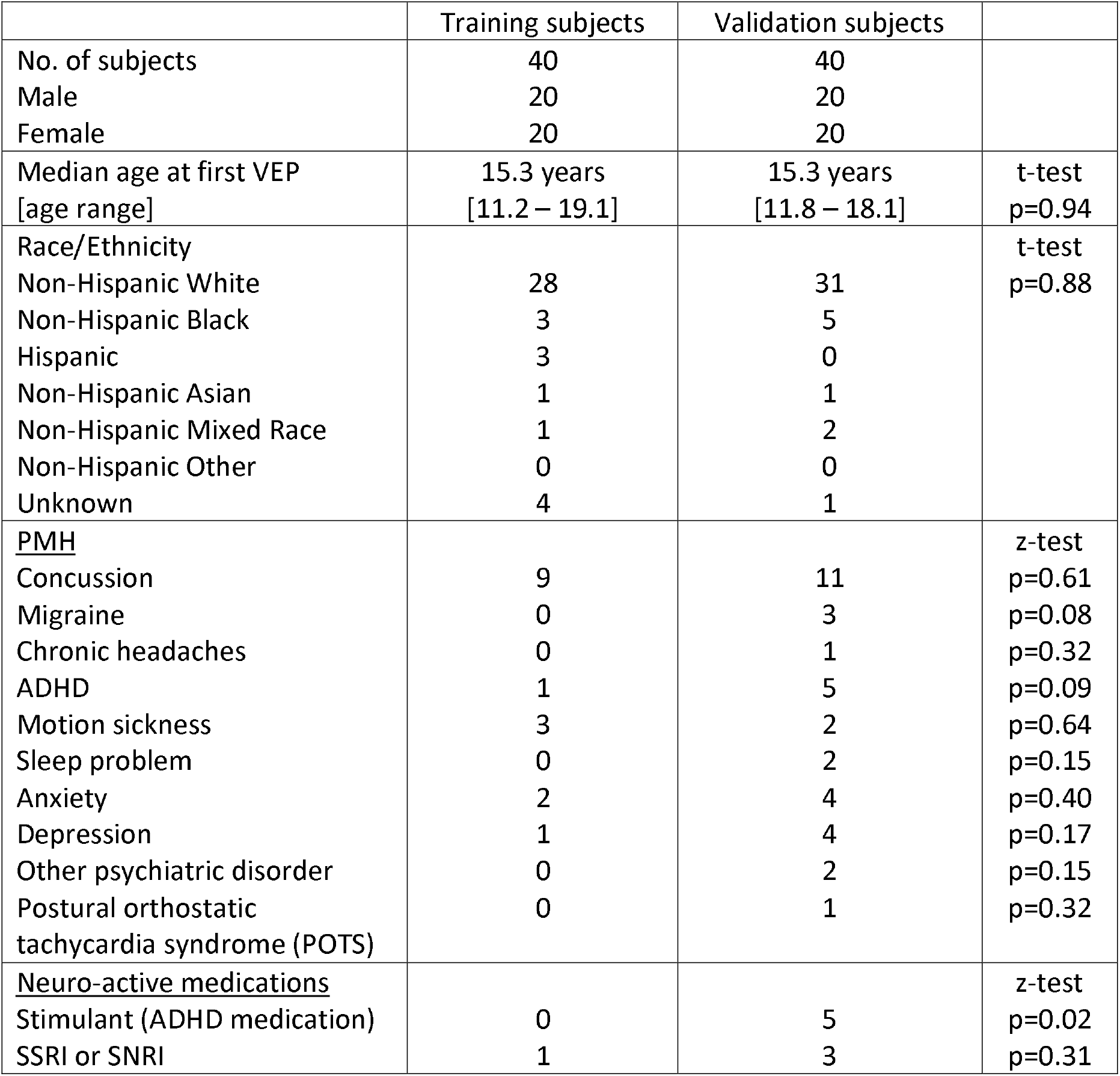
Subject demographics for the training and validation subject groups used to generate and validate the PCA model, respectively. No subjects reported a PMH of dyslexia, bipolar, drug/alcohol use disorder, autism, epilepsy, tic disorder, or amplified musculoskeletal pain syndrome (AMPS).

Measurements from the validation subjects were projected onto the PCA model generated from the training subjects. An example of a validation subject having their VEP projected upon the PCA model with resultant coefficients for the 7 PCs is shown in Figure 1b. If the PCA model is generalizable, it should be able to capture variance in the validation dataset similar to its performance in the training dataset. We found that the 7 PCs of the model accounted for 90.5% of the variance in the validation dataset.

As a second test of generalizability, we considered that the distributions of PC coefficients derived using the PCA model in the validation dataset should be similar to the distributions seen in the training dataset. We examined these distributions for each PC and found substantial overlap of the PC coefficients derived from the training (black) and validation (blue) datasets (Figure 3a). Comparison of the mean and distribution of PC coefficients between the datasets revealed no significant differences (Figure 3a, p>0.3 for all PCs; KS test). There was also no difference between training and validation datasets in the 7 PC multidimensional space (ANOVA F_(1,78)_=0.03, p=0.86).

**Figure 3:**
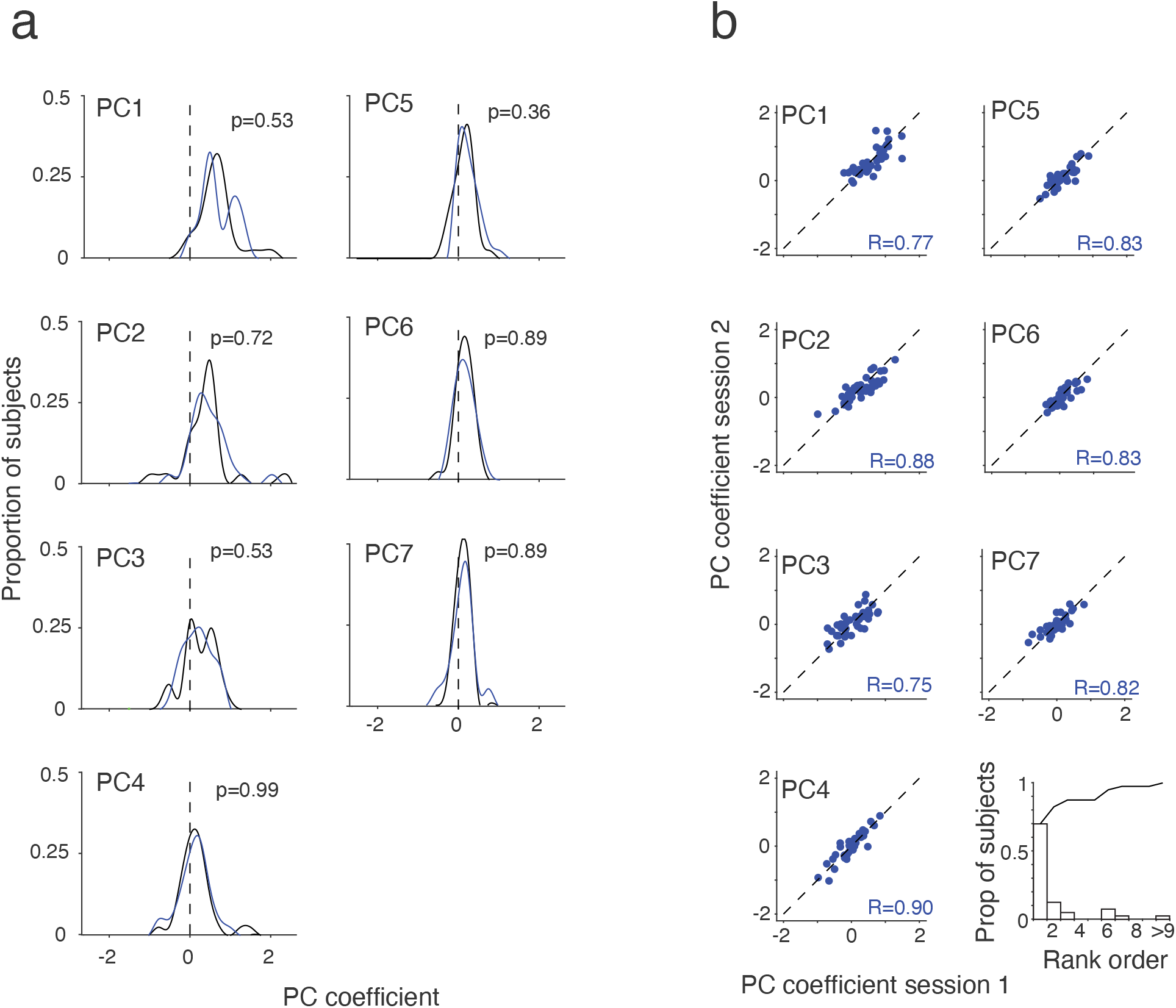
Generalizability and test-retest reliability of the PCA Model. (a) PC coefficent comparison of 40 training subjects (black) and 40 validation subjects (blue) showed no significant differences in the mean or distribution of individual PCs (KS test p>0.3 for all PCs) or the multidimensional space across all PCs (ANOVA F(1,78)=0.03, p=0.87). (b) Scatter plots present the coefficients for each PC dervied from session 1 and session 2 for each subject. There was high correlation for these measurements between sessions across subjects (all R>0.7). The bottom right panel shows the proportion of subjects with the rank score for a subject’s Euclidean difference between session 1 and session 2 compared to session 2 of all validation subjects. Black line shows the cumulative proportion of subjects.

### Coefficients derived using PCA model have good within subject re-test reliability

The PCA model was created using the validation dataset. Specifically, the mean age and mean prVEP was calculated from two separate sessions for each subject. We next asked if the PC coefficients obtained using this model were similar within subject for the two testing sessions. For each PC, we examined the correlation of coefficients across subjects for data from the first and second testing session (Figure 3b). The PCA model demonstrated high retest reliability across sessions (Pearson correlation coefficient, R ≥0.75 for all PCs). We examined the correlation between session 1 and session 2 coefficients across the first 40 components of the PCA model (i.e., extending beyond the first 7 PCs that we retained) (Figure S2a). The intra-subject reliability of the coefficients of the prVEP declines across the components, indicating that the higher dimensions of the model likely reflect non-reproducible noise in the measurement. Consequently, expressing prVEPs by projection onto the first 7 PCs of the PCA model constitutes a noise-reduction technique.

The similarity of any two prVEPs may be expressed by projecting the waveforms upon the PCA model and measuring the Euclidean distance between the two sets of 7 PC coefficients. The distance between session 1 and session 2 was calculated within the 7 PC dimensions for all subjects to determine if a subject could be correctly matched between their first and second sessions. This would indicate very high intra-subject compared to inter-subject reliability and indicate stability of VEP signal for a single subject across sessions. Rank order for the subject’s own match between session 1 and session 2, compared to combinations with all other subjects is shown (Figure 3b, lower right). In 70% of the subjects, the best match to their session 1 prVEP was their session 2 prVEP. In over 85% of the subjects, the session 2 prVEP was in the top 5 ranking of prVEPs that were similar to session 1, compared to 40 possible subject session 1 and session 2 combinations. We examined the ability to match the prVEP from a subject across sessions as a function of the number of PCs used to describe the data. Accuracy plateaued with inclusion of approximately 7 PCs further supporting the use of this dimensionality to describe the prVEP data (Figure S2b). We also looked at correlation between session 1 and session 2 for standard peak analysis metrics including the N75 and P100 peak latencies and the N75 – P100 peak-to-peak amplitudes. N75 and P100 peak latency showed good correlation (R=0.45, p=0.004; R=0.64, p<0.001, respectively), and the N75 – P100 peak-to-peak amplitude showed excellent correlation (R=0.92, p<0.001) across sessions (Figure S3a). Given the high test-retest reliability of the measurements, the mean prVEP was calculated across two sessions for each subject (when available) for the remaining analyses.

### Characteristics of the full cohort

The remaining analyses were performed on the full cohort of 155 subjects. Demographics are shown for the full cohort (Table 2). Two subjects had a past medical history of a ‘lazy eye’, two subjects had a history of eye patching, and one subject had a history of strabismus. Exclusion of these 5 subjects did not significantly change the outcome of the results. Seventeen (11%) of subjects reported taking neuroactive medications including stimulants for ADHD (9), selective serotonin reuptake inhibitors or serotonin and norepinephrine reuptake inhibitors (7), beta blockers (1), and antipsychotics (1). The other 34 subjects who reported medication use included as needed asthma inhalers, birth control, and allergy medications. Overall PCSI scores were low (median=3.5 out of a possible total score of 126), and there was not a significant difference in the distribution of PCSI scores for subjects with versus without a remote history of a concussion (p=0.53, KS test).

**Table 2:**
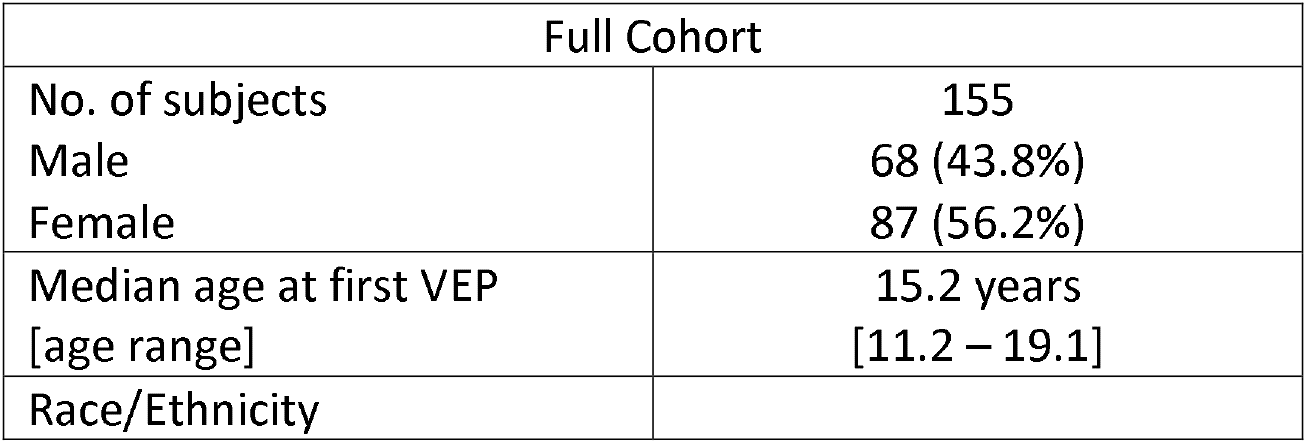
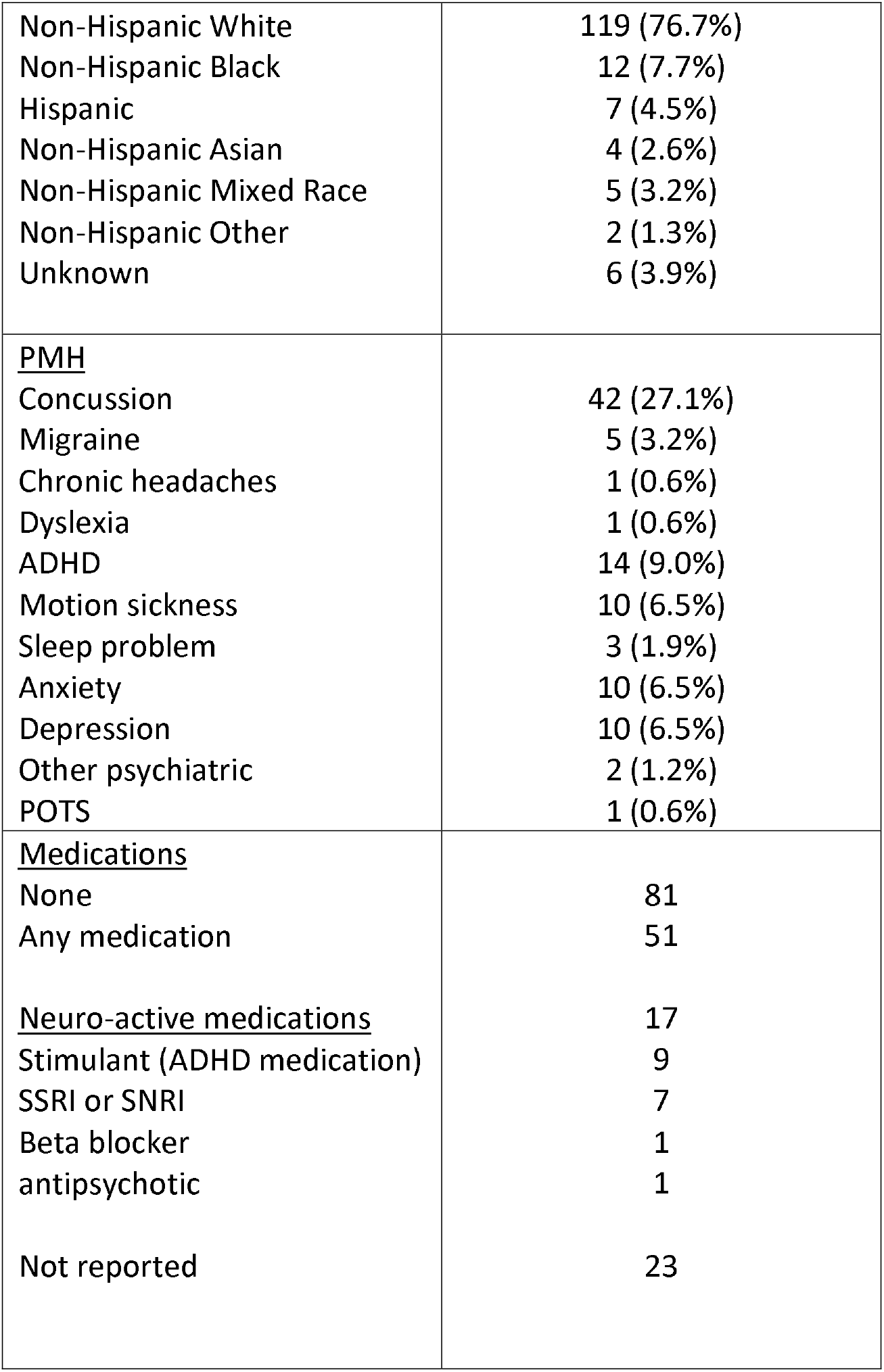
Subject demographics of the full pediatric cohort. No subjects reported a PMH of bipolar, drug/alcohol use disorder, autism, epilepsy, tic disorder, or AMPS.

### There was an effect of sex on prVEP in the PCA model

A linear regression model was generated to examine if the 7 PCs varied based on the sex of a subject. We did not observe a significant difference between males and females across all 155 subject (F_(7,147)_=1.69, p=0.12). However, when subjects on neuro-active medication were excluded (8 females and 9 males), there was a significant effect of sex on the PCA model (F_(7,130)_=2.31, p=0.03). Coefficients that showed a significant difference between male and female subjects were PC1 (t_(1,130)_=-2.13, p=0.03) and PC6 (t_(1,130)_=-2.48, p=0.01). These differences capture an increased overall amplitude (PC1) and a narrower P100 peak (PC6) for females compared to males. Mean male and female prVEPs and the distribution of the 7PC coefficients with and without inclusion of subjects on neuro-active medications are shown (Figure S4).

### The PCA model captures variability throughout maturation in our pediatric cohort

A linear regression model was generated to examine the effect of the 7 PCs upon the age of a subject. There was a significant, omnibus effect of age upon the 7 PC coefficients (F_(7,147)_=4.37, p=0.0002). The PC coefficients that showed a significant correlation with age were PC2 (t_(1,147)_=- 4.06, p=8e^−5^) and PC3 (t_(1,147)_=-3.21, p=0.002). PC coefficients for the 7 PCs are shown as a function age (Figure S5). The prVEP from any one subject can be described as a point in the 7-dimensional PC space. We plotted the prVEPs for our subjects along the sub-set of two dimensions that showed a significant change with maturation (Figure 4a). A vector within the PC space describes the effect of age upon the coefficients. The PCA model may be used to synthesize representative prVEP waveforms that lie along this vector. The result is a synthetic waveform that expresses the central tendency of the prVEP corresponding to subjects of different ages (Figure 4b). These waveforms capture multiple, non-local changes as a function of increasing subject age, including a progressively smaller P100 amplitude, narrowing of the P100 peak, and an increasing N135 amplitude. In agreement with these findings, peak analysis metrics showed a negative correlation of the N75-P100 peak-to-peak amplitude with age (R=- 30, p=0.002), but there was no significant correlation between age and peak latency (Figure S3b). Removal of subjects on neuro-active medications did not have a significant impact on the outcome of the PCA model across age.

**Figure 4:**
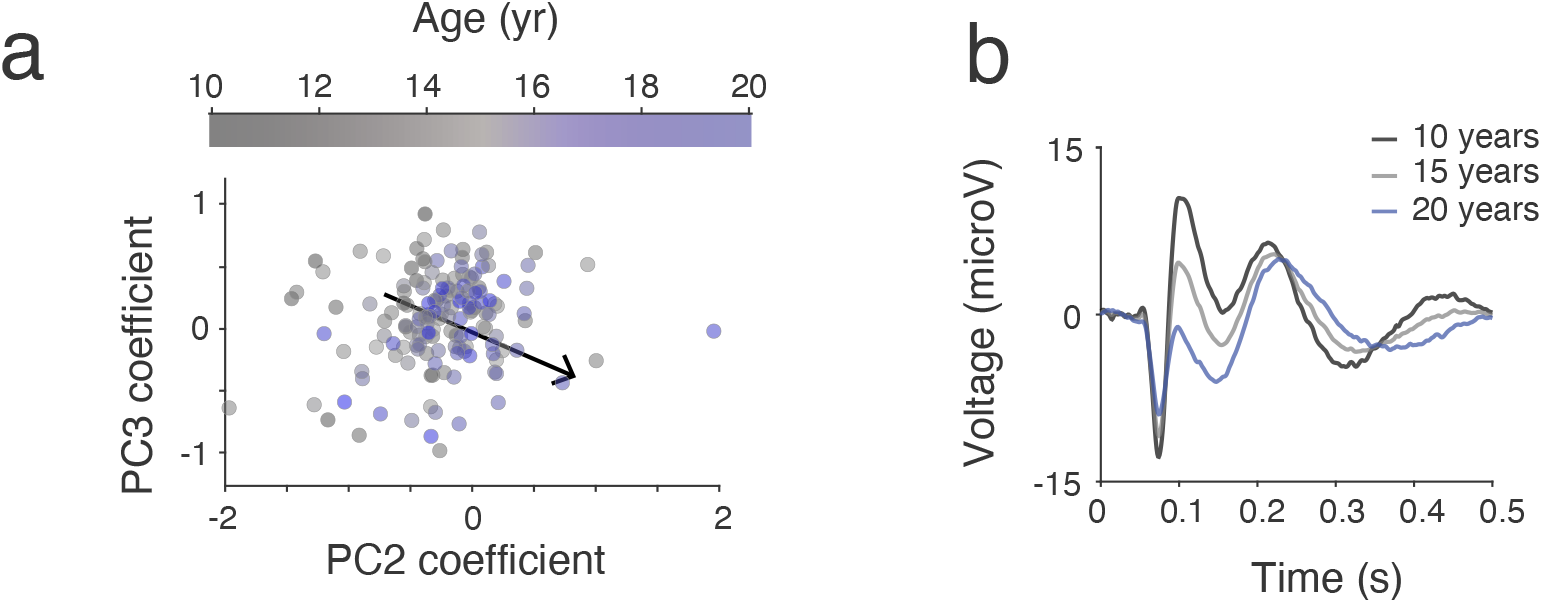
PC coefficients change systematically with age. (a) The response from each may be described by their PC2 and PC3 coefficients. These dimensions of the PCA model had a significant association with subject age. Plot points are colored from black to blue indicating increasing subject age, and the red arrow indicates the vector direction of the effect of subject age in this space. (b) Simulated VEPs for a 10-year old, 15-year old, and 20-year old subject.

We considered if a non-neural developmental change such as head circumference could account for age effects in the prVEP. We examined the correlation between age and head circumference in the 124 subjects with this measure and found a non-significant correlation (R=-0.04, p=0.66). There was no relationship between individual differences in head circumference and the set of 7 PC coefficients across subjects (F_(7,116)_=0.70, p=0.67). It seems therefore that variation in head circumference in this cohort has minimal effect upon our prVEP measures.

## DISCUSSION

We found that a PCA model using 7 PCs accounted for over 90% of the inter-subject variability of prVEP across 40 pediatric subjects for both a training and validation dataset. We demonstrate that this PCA model is a generalizable and reliable means of analyzing prVEP in a large pediatric sample. PCA offers a method of adjusting for differences across maturation to remove this confounding variability, which is important for the interpretation of VEP measurements in developing youth. For these reasons, PCA offers a promising complement to standard peak analysis for interpretation of VEP.

### Comparison to previous studies

Prior studies that have relied on peak analysis have generally found decreasing peak amplitude and decreasing peak latency with maturation, though some conflicting results have been reported. Snyder and colleagues (1981) found that the amplitudes of the P50, N64, P100, and N150 peaks decreased with increasing age during adolescence^12^. In related work, Shearer and Dustman (1980) found that latencies of these peaks gradually increased over time^13^. Wright and colleagues (1985) found overall amplitudes were higher in their youngest age group (10-19 years) compared to the older age groups, and found no differences in peak latencies across age^11^. One limitation of this study was that prVEPs were averaged across subjects in 10-year age groups, which would have failed to capture differences of maturation during adolescence. Emmerson-Hanover and colleagues (1994), who examined a large cohort of 406 subjects ages 6 to 80 years, found somewhat different results. They observed that the P50-N70 amplitude increased until about age 13 years and then decreased, while the N70-P100 amplitude decreased with maturation^10^. They also found the P50, N70, and P100 peak latencies decreased during maturation^10^. Brecelj and colleagues (2002) also found in children 7-18 years old that P100 peak latency decreased with maturation^18^. Allison and colleagues (1983) similarly found the VEP P100 peak latency decreased between 4-19 years of age, although the peak latency of other peaks and peak amplitude were not reported^9^. We also observed that prVEP amplitude decreases with maturation using both the PCA model and peak analysis (see Figure 4 and Figure S3b). Our PCA model also revealed a narrowing of the P100 peak with maturation that cannot be easily captured by standard peak analysis (see Figure 4b). Multiple PCs show a prominent change in the vicinity of the P100 peak. This reflects that there is substantial individual variation in the shape and the amplitude of the waveform in this temporal window. Age-related variability was predominantly described by PC2 and PC3. PrVEP changes with age in youth may be due to maturation of neural circuits within the visual pathway^18^. Head circumference did not appear to account for the differences we observed across age, though we cannot rule out the possibility that other physical characteristics such as skull thickness^19^ or age-related closing of cranial sutures^20^ could have played a role. Of note, our model of age-related effects over adolescence may not generalize to other age groups. The use of PCA in younger and older age groups would be an interesting expansion of the work presented here.

We find good generalizability of the PCA model between training and validation subjects. Indeed, there were no differences in PC coefficients for the training and validation datasets even though there were a few subjects in the validation set on stimulant medications for ADHD. We also find good intra-subject reliability for prVEPs in our pediatric cohort using both the PCA model and standard peak analysis metrics. An advantage with the PCA model is it includes 7-dimensions for comparison, which we show improves the performance of matching a subject’s session 1 to their own session 2 (Figure S2b). We are unaware of a similar prior measurement in children, although the prVEP has also been found to have good reliability in adults^21–23^. Very few studies have examined the stability of VEP signals across sessions in the pediatric population. Schellberg and colleagues (1987) found substantial intra-subject variation of flash VEP between sessions in 26 children 10 to 13 years old spaced 10 months apart on average. The variability they reported was higher than what was observed in adult studies^14^. Their use of flash VEP, which can show higher intra-subject variability^15^, may at least partially account for the high intra-subject variability. Differences may also be in part to reliance on peak analysis compared to PCA.

Prior work indicates that females tend to have increased peak amplitudes and shorter peak latencies compared to males^24–26^. The PCA model revealed a broader and lower amplitude P100 peak in males, which is consistent with these prior results while providing a richer account of the differences in VEP waveforms between sexes. This effect achieved significance when subjects on neuro-active medications were excluded from the analysis (figure S4).

### Global temporal analysis of VEP

Standard peak analysis of VEP remains an important tool for measuring the integrity of the visual system in many neurologic and ophthalmologic conditions. However, a focus upon the peak amplitude and peak latency in the prVEP necessarily limits the ability of an analysis to detect subtle, non-local changes in the shape of the prVEP waveform^22^. Sarnthein and colleagues (2009) used a combination of a metric of VEP shape and traditional peak analysis to address this limitation. They similarly demonstrated high test-retest reliability of prVEPs over 8 months in 10 healthy adult women using a combination of N75 and P100 peak-to-peak amplitude and peak time with pairwise regression of VEP waveforms to account for VEP shape^22^. Here we describe a method using PCA that takes this one step further in eliminating focus on predefined peaks and quantifying instead the most informative individual variability in the temporal waveform.

Though there are many advantages, the PCA method has the limitation that the components are not temporally constrained so they cannot be as easily be linked to discrete temporal physiologic events^27^. Many studies have focused on linking the N75, P100, and N135 peaks of the prVEP to different stages of processing in the visual hierarchy^28^. Compromise between the unconstrained decomposition of PCA and a rigid focus on individual peaks may be found in the use of informed basis functions, as has been used in the spatial analysis of fMRI^29^ and EEG^30^ data. In such an approach, the prVEP is projected onto a set of components that are crafted to reflect both temporally local features and global variation. The development and use of informed basis sets offers a promising direction for future prVEP analysis.

While PCA has been applied to spatial localization in multi-focal VEP studies^27,31^, we are unable to find prior examples of PCA or other dimensionality reduction approaches being used to characterize individual differences in the timeseries of prVEP or other EEG data. Indeed, there have been calls to increase the use of model-based inference in EEG^32^. We have demonstrated that modeling the global temporal variability of prVEP using PCA is highly generalizable and repeatable in a pediatric cohort. These results are promising for the application of this approach to the study longitudinal pediatric data. Further, our modeling approach may be used to capture and remove variability associated with maturation. This offers a means of addressing confounds of unmatched age differences in the populations, and testing for the subtle effects of a disease upon maturation. Finally, this model has the potential to identify differences between normal and pathologic groups that are not captured by focusing on specific pre-defined peaks.

## CONCLUSION

PCA is a highly repeatable and generalizable method of analyzing prVEP data and offers a useful means of addressing variability during maturation in youth. These features are especially advantageous for longitudinal study designs. PCA offers a complimentary approach to standard peak analysis that can account for the global temporal variability in the prVEP waveform, which warrants further study in both healthy and diseased states.

## Acknowledgements

The authors would like to thank Fairuz Mohammed, Anne Mozel, and Olivia Podolak for their support in collecting and organizing the data in these experiments. The authors wish to thank the two anonymous reviewers, who provided patient, helpful, and constructive advice that markedly improved the manuscript and saved us from errors.

## Abbreviations

ADHD: Attention Deficit Hyperactivity Disorder
AMPS: Amplified Pain Syndrome
fMRI: functional magnetic resonance imaging
EEG: electroencephalogram
PCA: Principal Component Analysis
PC: Principal Component
PMH: Past Medical History
POTS: Postural orthostatic tachycardia syndrome
prVEP: Pattern reversal visual evoked potential
SNRI: Serotonin and norepinephrine reuptake inhibitor
SSRI: Selective serotonin reuptake inhibitor
SVD: Singular value decomposition

