## Supplemental figures for "Developmental effects on pattern visual evoked potentials characterized by principal component analysis"

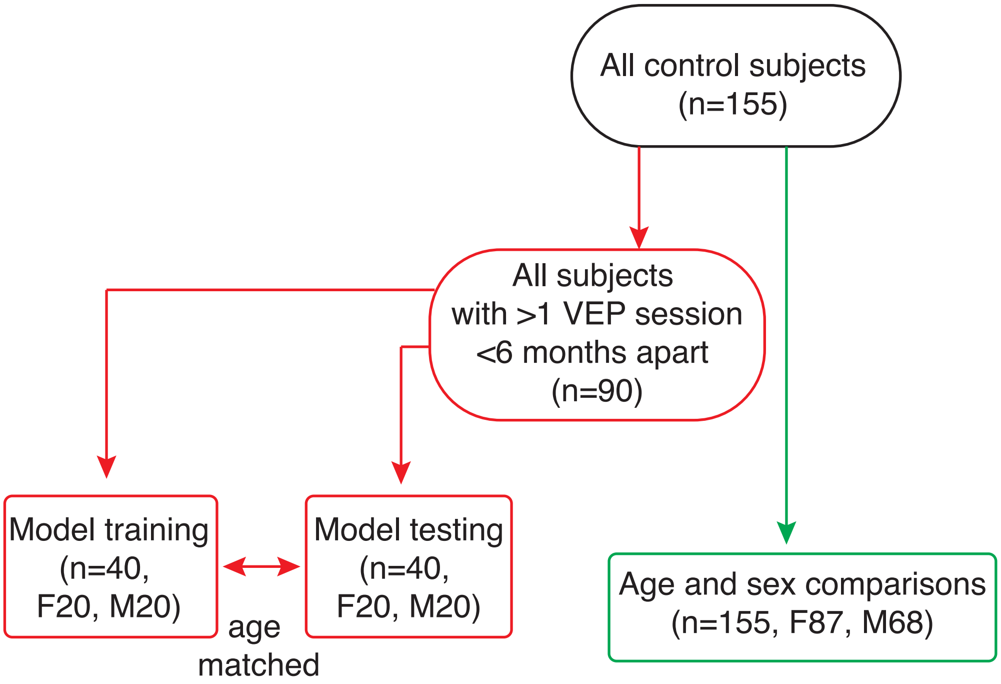


Figure S1: Subject flow chart. Flow chart illustrates subjects used in Principal Component Analysis (PCA) model training and test (red), and age and sex comparisons (green). F stands for female, M stands for male, n is number of subjects.

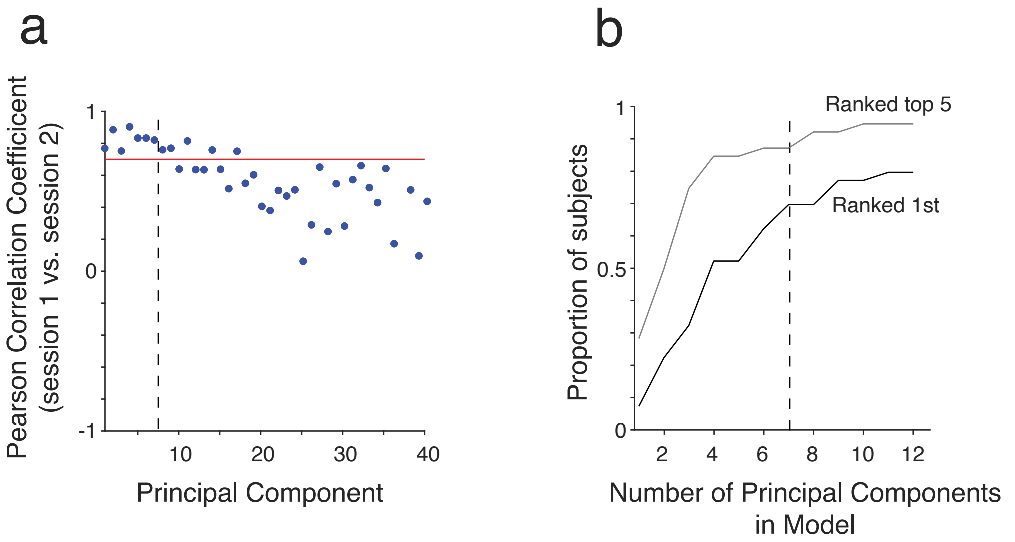


Figure S2: PCA Model reliability by PC. (a) Pearson correlation coefficent for PCs 1 to 40 (in order of decreasing variability accounted for in the data) are shown for the test data set. Red line indicates R of 0.7 for high reliability. Dotted line separates the first 7 PCs. (b) Proportion of subjects whose own session 2 ranked first (black) and in the top 5 (gray) in the closest Euclidean distance compared to session 1 as a function of the number of PCs included in the model. Dotted line shows the model with 7 PCs.

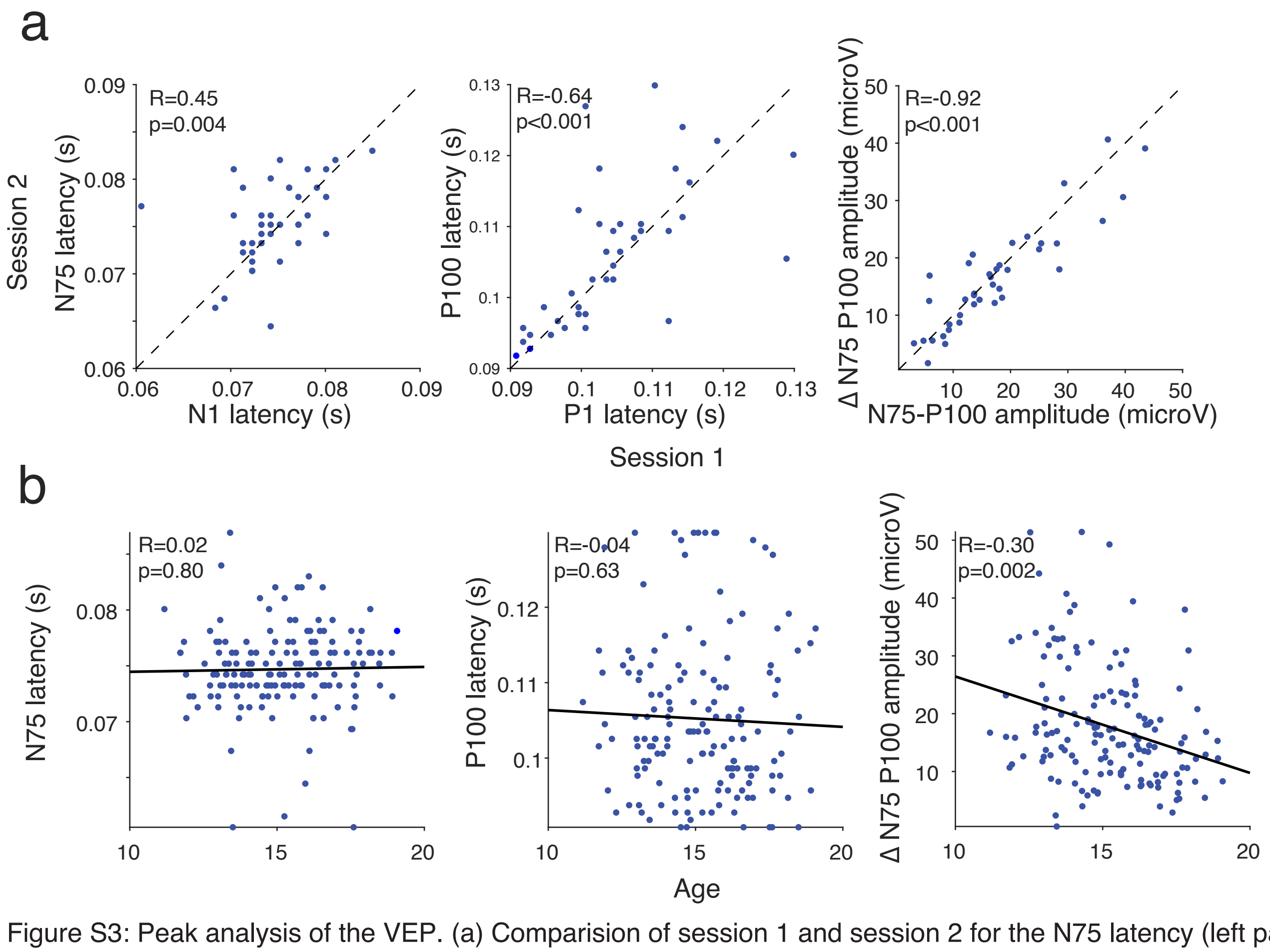


Figure S3: Peak analysis of the VEP. (a) Comparision of session 1 and session 2 for the N75 latency (left panel), P100 latency (middle panel), Δ N75 P100 peak-to-peak amplitude (right panel) for the 40 validation subjects. (b) Correlation between subject age and N75 latency (left panel), P100 latency (middle panel), and Δ N75 P100 peak-to-peak amplitude (right panel) for all 155 subjects. Pearson correlation coefficients with p values are shown.

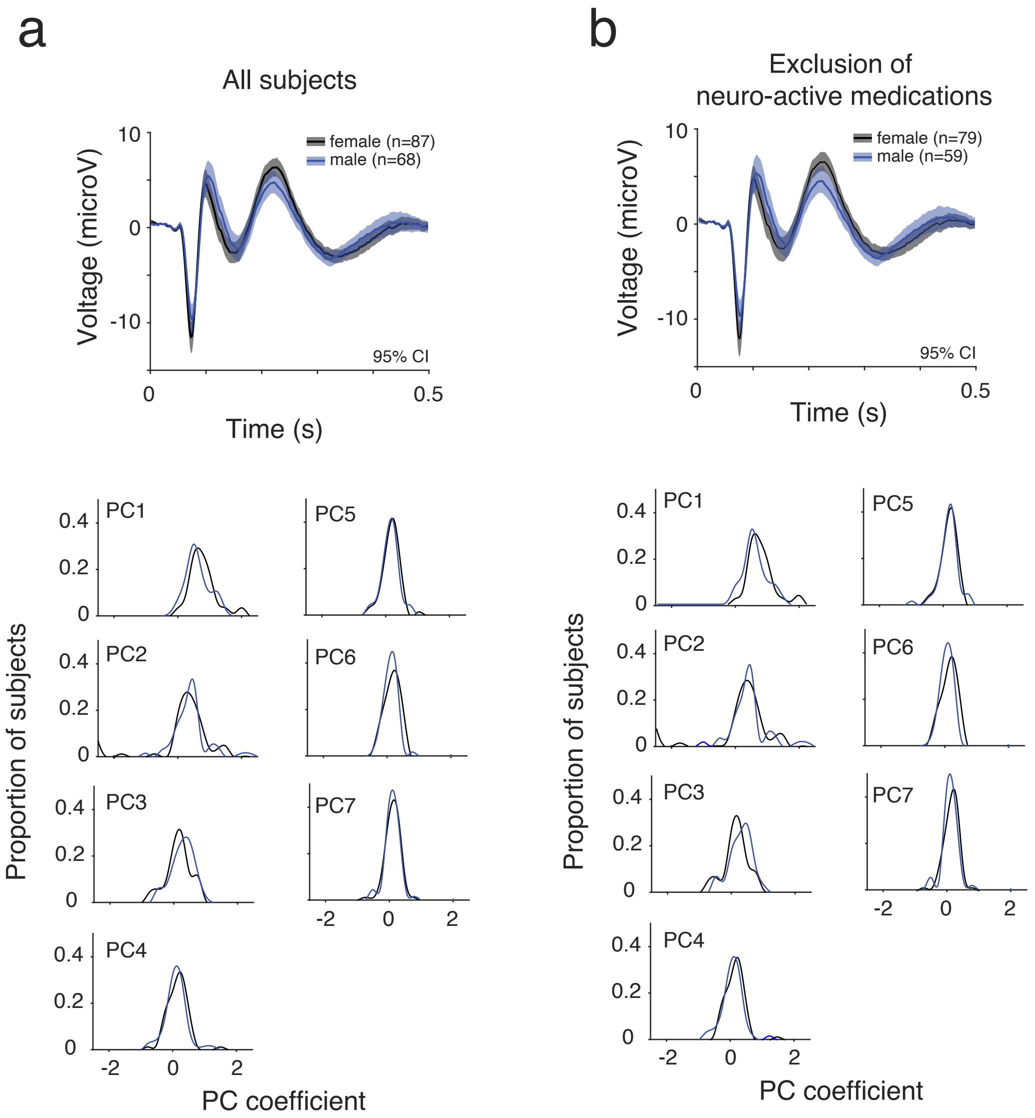


Figure S4: Male and female comparison. Mean VEP for males and females (top panel) and PC coefficients (bottom panel) for (a) the full cohort (n=155), and (b) only subjects who were not on neuroactive medications (n=138).

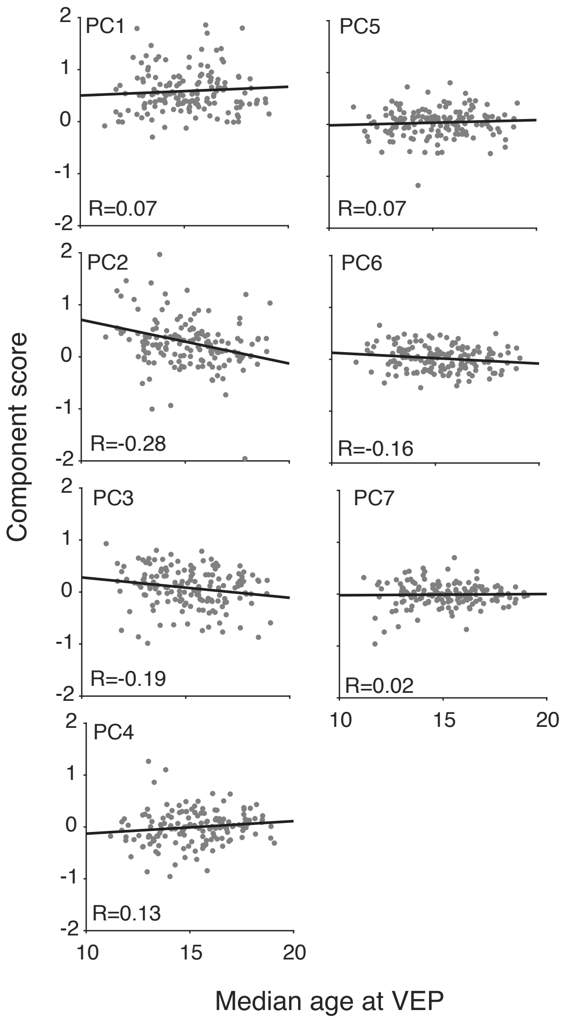


Figure S5: Unadjusted PC coefficients ploted as a function of age for the 7 PCs of the PCA model. Black line represents linear least sqaures fit.
